# qMAP enabled microanatomical mapping of human skin aging

**DOI:** 10.1101/2024.04.03.588011

**Authors:** Kyu Sang Han, Inbal B. Sander, Jacqueline Kumer, Eric Resnick, Clare Booth, Guoqing Cheng, Yebin Im, Bartholomew Starich, Ashley L. Kiemen, Jude M. Phillip, Sashank Reddy, Corrine E. Joshu, Joel C. Sunshine, Jeremy D. Walston, Denis Wirtz, Pei-Hsun Wu

## Abstract

Aging is a major driver of diseases in humans. Identifying features associated with aging is essential for designing robust intervention strategies and discovering novel biomarkers of aging. Extensive studies at both the molecular and organ/whole-body physiological scales have helped determined features associated with aging. However, the lack of meso-scale studies, particularly at the tissue level, limits the ability to translate findings made at molecular scale to impaired tissue functions associated with aging. In this work, we established a tissue image analysis workflow - quantitative micro-anatomical phenotyping (qMAP) - that leverages deep learning and machine vision to fully label tissue and cellular compartments in tissue sections. The fully mapped tissue images address the challenges of finding an interpretable feature set to quantitatively profile age-related microanatomic changes. We optimized qMAP for skin tissues and applied it to a cohort of 99 donors aged 14 to 92. We extracted 914 microanatomic features and found that a broad spectrum of these features, represented by 10 cores processes, are strongly associated with aging. Our analysis shows that microanatomical features of the skin can predict aging with a mean absolute error (MAE) of 7.7 years, comparable to state-of-the-art epigenetic clocks. Our study demonstrates that tissue-level architectural changes are strongly associated with aging and represent a novel category of aging biomarkers that complement molecular markers. Our results highlight the complex and underexplored multi-scale relationship between molecular and tissue microanatomic scales.

## Introduction

Aging is a complex process marked by the gradual decline of physiological functions and deterioration across various organs and tissues^1^. Biological age, as opposed to chronological age, is a key risk factor for numerous diseases and health conditions that impact overall health and lifespan of individuals^2^. Recent advances in geroscience have revealed connections between seemingly unrelated diseases, highlighting the central role of aging in their onset and progression^3^. It is imperative to understand the natural aging process and the alterations occurring in biological systems over time to pinpoint novel therapeutic avenues and effective biomarkers for aging. Identifying reliable and robust biomarkers of biological aging is critical for accurately stratifying the risk of individuals to design anti-aging interventions and novel therapeutic strategies^4^.

As individuals age, they tend to experience progressive dysfunctions, including those of the cardiovascular, musculoskeletal, and respiratory systems^5–7^. Extensive research efforts have revealed features of biological systems over a wide range of length scales, from the molecular scale to the organ scale, that change over time^8–10^. At the molecular level, advances in omics techniques have facilitated the development of aging biomarkers^11,12^ in genetics^13,14^, epigenetics^15–17^, transcriptomics^18,19^, proteomics^20–23^, and metabolomics^19^. At the organ level, aging is associated with changes in blood biochemistry^23^, immune cells composition^24–26^, and morphology of organs^5–7^.

As the functions of biological systems emerge from the integration of multiple scales, including the molecular, cellular, tissue, and physiological levels, exploring the association and utility of aging at the cellular and tissue scales is crucial^27^. These scales are critical for bridging changes at the molecular scales to the physiological scales, thereby facilitating a deeper understanding of the aging process and promoting the delineation of chronological and biological aging biomarkers. Examining histological tissue sections is one of the primary approaches to simultaneously study biological systems at both the cellular and tissue scales^28–32^. However, performing effective quantitative analysis on tissue sections has been a challenging task due to the spatial complexity and variability of tissue structures, as well as the limitations of analytical techniques^33^. These challenges hinder the utility of tissue architectural information as potential biomarkers of aging^28^.

In this study, we developed a deep-learning-based tissue image analysis pipeline named quantitative micro-anatomical phenotyping (qMAP). This pipeline quantitatively profiles microanatomical tissue and cellular features in histological sections stained with hematoxylin and eosin (H&E). qMAP accurately labels distinct tissue compartments and individual cells from whole-slide tissue images using deep learning frameworks. We implemented qMAP to quantitatively profile microanatomic changes in human skin of the back and explore the potential use of tissue architectural features as aging biomarkers.

## Results

### Overall workflow of human skin tissue analysis

To investigate age-related changes in non-diseased skin, we obtained 379 formalin-fixed, paraffin-embedded (FFPE) skin biopsy specimens from patients at the Johns Hopkins Hospital collected between 2015 and 2021. These samples comprised histologically normal adjacent tissues from excision specimen of patients diagnosed with various conditions, such as melanoma, seborrheic keratosis, epidermal cysts, fibrosis, basal cell carcinoma, and squamous cell carcinoma. A board-certified dermatopathologist reviewed all H&E slides for each tissue specimen and identified 167 patient cases with substantial healthy margin, defined by the absence of disease or other damage, such as solar elastosis, excessive inflammation, or cicatrix (**Figure 1a and Supplementary Figure 1**). Among these, the majority (N=130) were skin tissue blocks from the back, with the remainder from the abdomen, chest, head, neck, arms, and legs. Out of the 130 back tissues, 99 originated from Caucasian donors; with 56 males and 43 females. The samples were split into three age groups: young (age<30), middle-aged (30≤age≤60), and old (60<age). Of the 99 donors, 17 were young, 41 middle-aged, and 41 old (**Supplementary Table 1**).

**Fig. 1:**
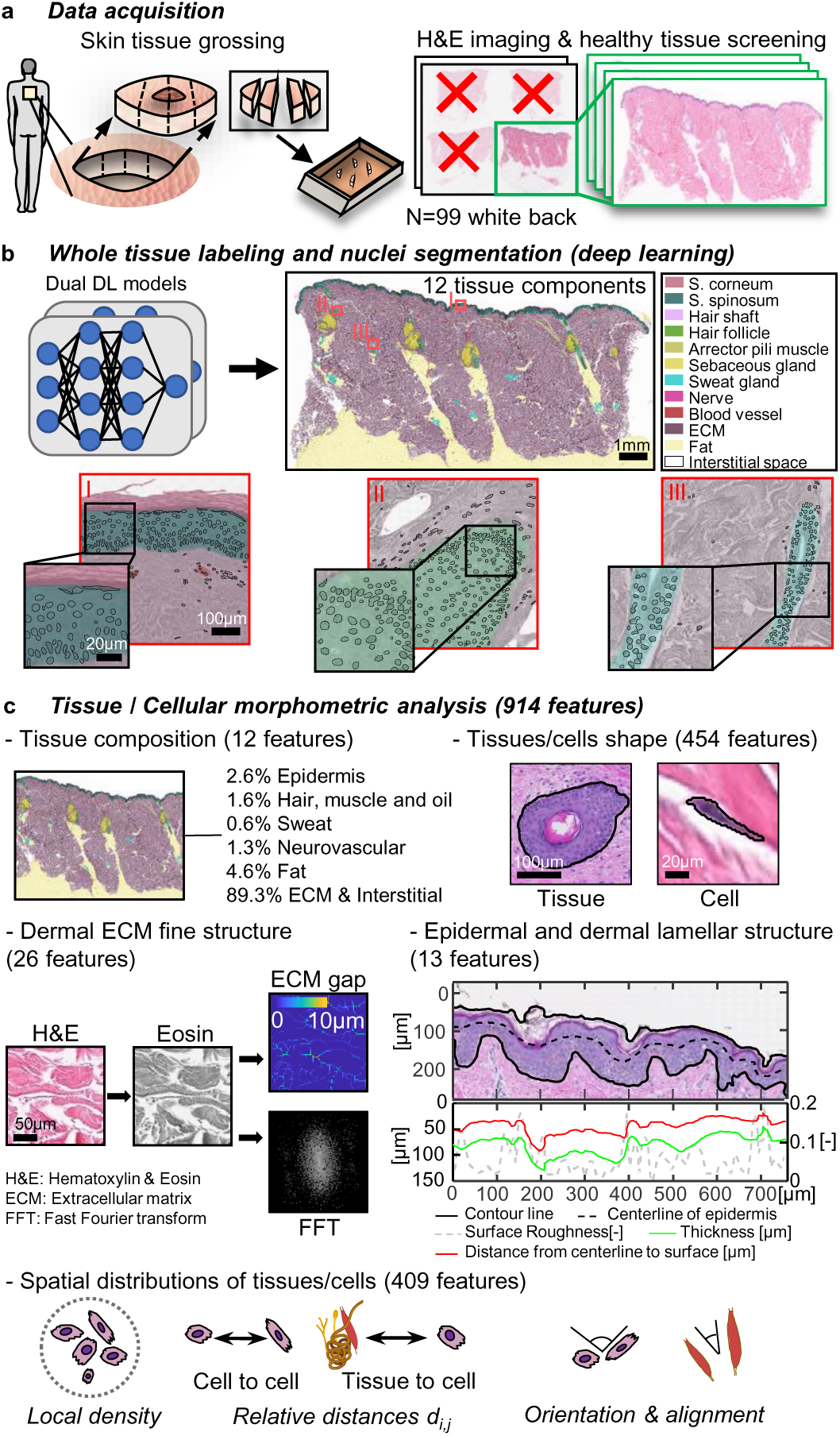
Overall workflow of human skin tissue analysis. **a.** Skin excisional sample typically with an elliptical shape is cut into multiple pieces before embedding into a paraffin block. The skin pieces with abnormal tissue architecture (such as lesion, sun damage, …etc.) are excluded for further analysis through examining the image of H&E-stained tissue section. **b**. Whole slide H&E images are labeled with tissue components automatically using a trained CNN with DeepLab V3+ framework. Cell nuclei within each labeled tissue component are segmented using HoVerNet. Scale bar, 1mm in tissue, 100µm and 20µm in zoom-in **c**. Five groups of morphometric features with a total of 914 features are established to characterize the tissue and cellular architecture of skin. Tissue scale bar, 100µm. Cell scale bar, 20µm. H&E scale bar, 50µm.

To comprehensively analyze the micro-architecture of the skin and its constituent cells, we spatially mapped the architecture of the tissue and cell components in these tissue specimens. We identified twelve primary microanatomical compartments distinguishable from H&E images: 1. Stratum corneum, 2. Stratum spinosum, 3. Hair shaft, 4. Hair follicle, 5. Smooth muscle, 6. Sebaceous gland, 7. Eccrine coils (sweat gland), 8. Nerve, 9. Vasculature, 10. Extracellular matrix (ECM), 11. Fat, and 12. Interstitial space (**Figure 1b** and **Supplementary Figure 2**). We established a deep learning-based workflow that precisely delineated these tissue microanatomical compartments (i.e. tissue compartment labeling) and the nuclei of the cells contained in these compartments. This was achieved by combining the DeepLabV3+ tissue labeling pipeline^34^ with HoVerNet for nuclei segmentation^35^ (**Figure 1b**). To train and validate our deep learning model for tissue compartment labeling, we performed manual annotations on 50 instances of each tissue microanatomical compartment in 20 skin samples. Out of these samples, 15 were allocated for training, and the model’s performance was assessed on the remaining 5 unseen tissue images. The trained model achieved a precision of 96.6% and an average recall of 93.9% across all tissue types (**Supplementary Figure 3**).

To segment the nuclei in skin tissues, we first tested the performance of the pretrained HoVerNet model trained on CoNSeP pancreatic tissue images^35^. The pretrained model achieved a DICE score of 0.66, DICE is defined as 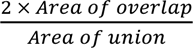 when the predicted nuclei are compared to the manually annotated nuclei. To improve nuclei segmentation, we developed a semi-supervised learning workflow (**Supplementary Figure 4**) based on the model training strategy^36^ that does not require additional manual annotations or curation to re-train the model on the skin tissue images. Our results showed that this semi-supervised model substantially improved nuclei segmentation on skin with a DICE score of 0.73 (**Supplementary Figure 4**).

The quantitative micro-anatomical phenotyping (qMAP) workflow begins with precise segmentation of both tissue and nuclei based on H&E-stained tissue slide images. The workflow is followed by the extraction of morphological features and their spatial information.

### Establishing a robust tissue and cell morphometric analysis

To quantitatively assess the association between tissue micro-architecture and cellular content within these skin tissues, we developed a panel consisting of 914 morphological features categorized into five groups: 1) tissue composition, 2) tissue and cell morphology, 3) dermal ECM structure, 4) epidermal and dermal lamellar structure, and 5) spatial distributions of tissues/cells (**Figure 1c**). The detailed features list is found in **Supplementary Table 2**. Of note, our tissue analysis focuses on the epidermis and dermis, and does not include the subcutaneous tissue.

Qualitative assessment of the skin images showed changes in tissue architecture, such as dermal thinning and epidermal flattening, indicating that changes in tissue composition and architecture were associated with aging (**Figure 2a-b**). Quantitative analysis of skin tissue composition, based on the deep learning labeled tissue component map, indicated that the average human back skin was composed of approximately 89.3% ECM and interstitial space, 4.6% intradermal fat, 2.6% epidermis, 1.5% pilosebaceous unit (including the hair shaft, hair follicle, sebaceous gland, and arrector pili muscle), 1.3% neurovascular bundle (consisting of nerve and blood vessel), and 0.6% sweat gland. (**Supplementary Figure 5**).

**Fig. 2:**
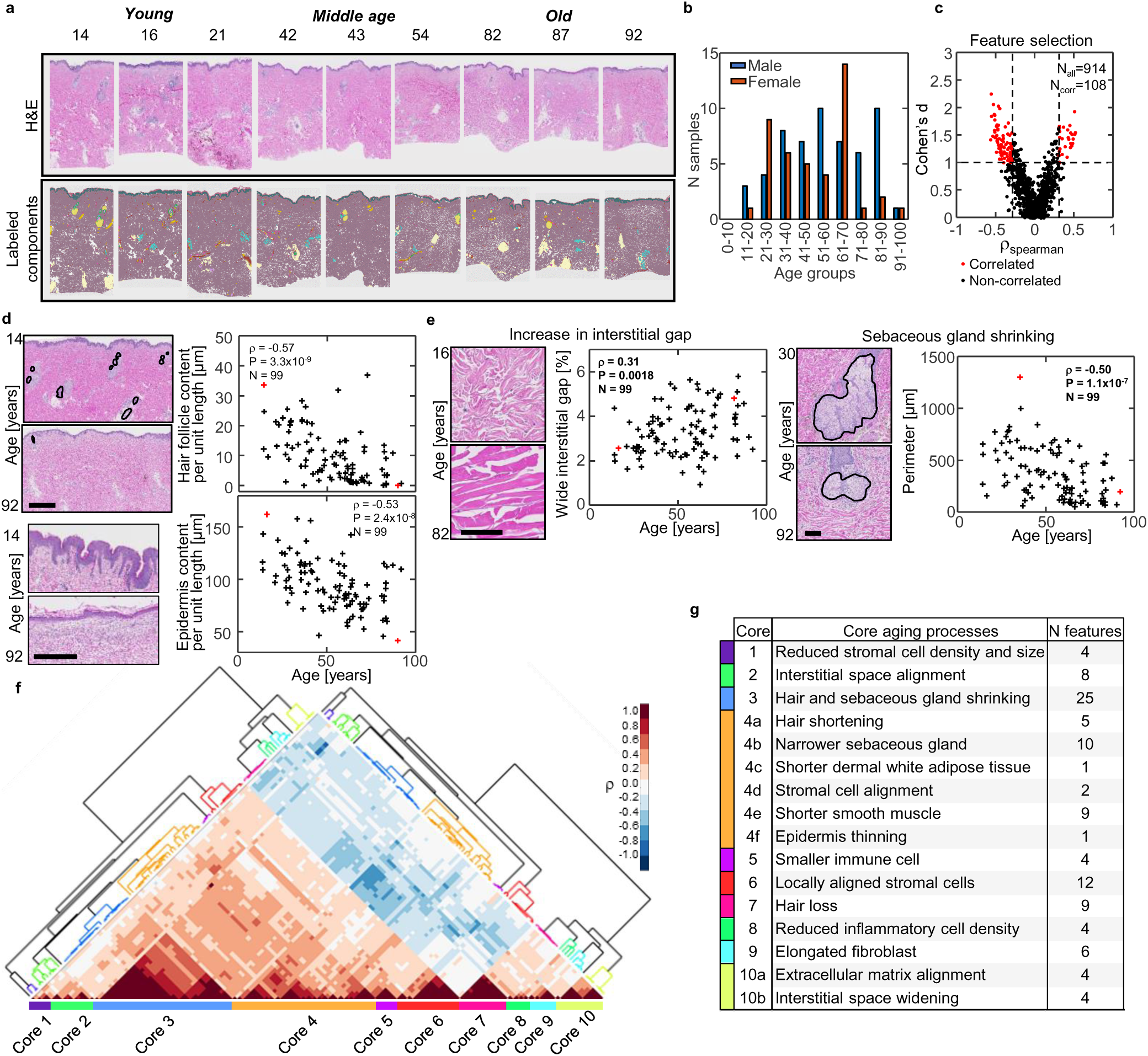
Core aging processes through extensive morphometric analysis. **a.** Representative image of H&E-stained tissue section from donors and at young, middle age and old (top). The corresponding tissue component maps labeled by the trained CNN (bottom). **b.** Histogram of age distribution of the sample cohort (56 male and 43 female) obtained from the back of skin. **c.** Age associated features are selected based on the spearman correlation coefficient with aging and Cohen’s d from comparing young (30 years old or younger) vs old (60 years old or older). A scatter plot showing each features spear correlation coefficient and Cohen’s d. A total of 108 features with Cohen’s d >1 and |spearman’s rho| >0.3 are identified as aging-associated features. **d.** Representative H&E images of skin and scatter plots showing the hair follicle content and epidermis content are among the identified aging-associated features. Hair follicle content and epidermis content are features known to be correlated with age. Scale bar, 1mm. **e.** Representative H&E images of skin and scatter plots show the increment of interstitial gap and shrinkage of the sebaceous gland, which are novel features of skin aging, among others. Scale bars, 100µm. Wide interstitial gap ratio is measure by ratio of the interstitial gap exceeding 4µm thickness among total by length. **f-g.** Hierarch clustering analysis groups the aging-associated features into ten distinct cores of the aging process using pair-wise correlation distance (**f**). Summary of general characteristics of each core based on the morphometric features of skin in each core (**g**). Cores 4 and 10 are further divided into subgroups to describe distinct characteristics of the assigned features better.

To identify potential aging-associated tissue changes in the microarchitecture of skin, we used a ranked-based Spearman correlation (r) of the 914 microarchitectural features with aging across all tissue samples (**Figure 2b**). In addition, we measured the Cohen’s distance (*d*), which describes the difference between two sample populations: samples from patients < 30 years and > 60 years old. We found that 108 features were significantly associated with aging with absolute values of r > 0.3, *p*-value < 0.05, and *d* > 1 (**Figure 2c, Supplementary Table 3**). Among these 108 features, we identified several features previously known to be associated with aging, such as epidermal thinning^37–41^ and loss of hair follicles and associated appendageal structures^42,43^, which validated our morphometrics analysis (**Figure 2d**). This approach allowed for the precise quantitation of those changes. The epidermal thickness decreased from 117µm to 100µm from young (less than 30 years old) to old (more than 60 years old) age groups. Across the tissue samples of all ages, this metric had a Spearman’s correlation of −0.36 and p-value of 2.4×10^−4^. The average reduction of epidermal thickness is 0.28 microns per year. Additionally, our results showed a significant decrease in the density of hair follicles, decreasing from 0.4% in young patient samples to 0.1% in old patient samples (Spearman’s correlation coefficient −0.55, p-value 5.11×10^−9^), and sebaceous glands, which decrease from 1.0% in young patient samples to 0.3% in old patient samples (Spearman’s correlation −0.42, p-value 1.3×10^−5^).

Besides these highly correlated features, we identified several features that had previously been thought to be associated with aging; however, the changes in these 806 unselected features were not significant. For example, dermal thickness decreased with age from 5.0mm to 4.8mm (Spearman’s correlation −0.16, p-value 0.10). The full list of unselected features is presented in **Supplementary Table 3**.

Importantly, we also established and identified several novel biomarkers of the aging skin, such as increased interstitial space, reduced sebaceous gland size, reduced hair density, and increased alignment of stromal cells to ECM components in the back skin (**Figure 2e**). We performed unsupervised hierarchical clustering analysis to group these 108 aging-associated features (**Figure 2f**). This analysis revealed that these features could be further grouped into ten distinct core aging processes, including: Core 1. Reduced stromal cell density and nuclei size, Core 2. Interstitial space alignment, Core 3. Hair and sebaceous gland shrinking, Core 4a. Hair shortening, Core 4b. Narrower sebaceous gland, Core 4c. Shorter dermal white adipose tissue, Core 4d. stromal cell alignment, Core 4e. Shorter smooth muscle, Core 4f. Epidermal thinning, Core 5. Smaller immune cells, Core 6. Stromal cell local alignment, Core 7. Hair loss, Core 8. Reduced inflammatory cell density, Core 9. Elongated fibroblasts, Core 10a. Extracellular matrix alignment, and Core 10b. Interstitial space widening (**Supplementary Table 4**). The analysis showed that core 3 (Hair and sebaceous gland shrinking) and core 4 (Shorter smooth muscle and stromal cell alignment) had the largest numbers of features, comprising 25 and 28 features, respectively (**Figure 2g**).

### Skin tissue microarchitecture as biomarker of aging

To assess whether the identified aging-related morphometric features of the skin might serve as predictive biomarkers of chronological age, we conducted analysis on the mean absolute error (MAE) associated with each individual feature predict age. This was carried out using leave-one-out validation combined with univariate generalized linear regression (GLM) (**Figure 3a**). Overall, the MAE from these individual features ranged from 13 to 29 years, with an average of 17 years and a standard deviation of 0.7 years (**Figure 3a and Supplementary Table 5**). The ten best predictors are listed in **Supplementary Table 5**. Among all aging-associated features, the reduction in blood vessel radius was the most accurate aging predictor with an MAE of 13.1 years (**Figure 3b**). This result may explain the reduced blood flow observed with aging^44^. Comparing the ten distinct core aging processes, we found that features from core 2 (Interstitial space alignment) and core 3 (Hair and sebaceous gland shrinking) had better univariate prediction with their univariate MAE of 14.9 years and 14.6 years, which were significantly lower than the MAE of other cores (**Figure 3c, Supplementary Table 4**).

**Fig. 3:**
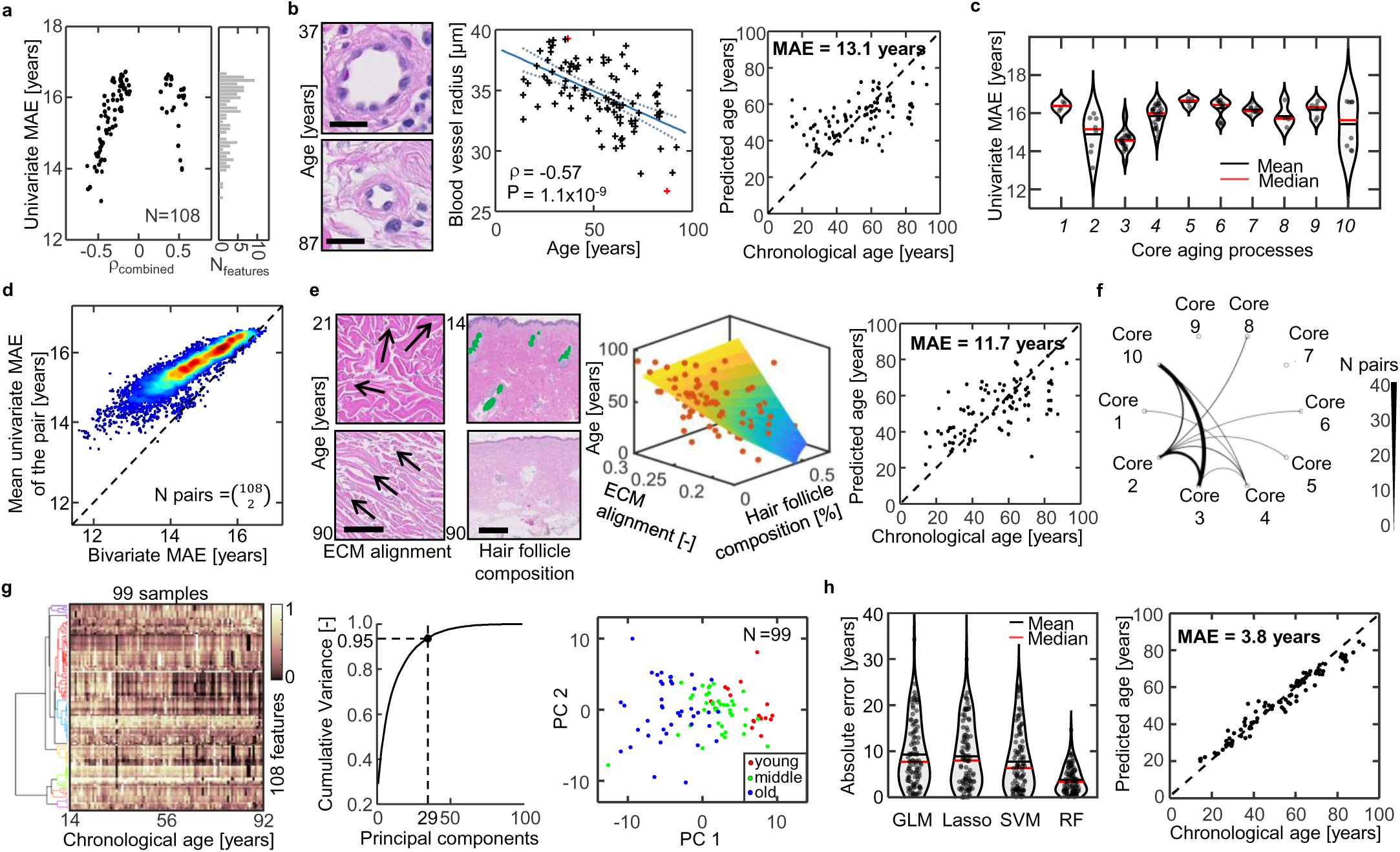
Morphometric features are predictive of chronological age. **a.** A scatter plot shows correlation coefficient with aging and mean absolute error of aging prediction for each individual aging associated feature. Mean absolute error is calculated from leave-one-out validation using univariate generalized linear regression model (GLM). **b.** The blood vessel radius is the best predictive feature with MAE of 13.1 years. Two representative H&E images from a young and an old donor are shown. Scale bar = 20µm. A scatter plot shows a negative correlation between blood vessel radius from each sample and their corresponding ages. Predicted age with blood vessel radius from leave-one-out validation and actual chronological ages for each sample is shown as a scatter plot. **c.** Violin plot shows MAE distribution for features in each core of aging process. Features from core 2 (interstitial space alignment) and core 3 (Hair and sebaceous gland shrinking) show the overall best performance in aging prediction as single variable among others. **d-f.** Aging prediction uses bivariate features. A scatter plot shows the MAEs from all pairs (N=5,778) of features using bivariate aging prediction with GLM compared to the average MAE of each pair with univariate prediction with GLM. The MAE is calculated using the leave-one-out validation. In most pairs, bivariate MAE is lower than the mean univariate MAE (**d**). The best predictive bivariate pair is ECM alignment and hair follicle composition with an MAE of 11.7 years. Representative H&E images of skin in young and old highlight decrement in ECM alignment and loss of hair follicles with aging. Scale bar, 100µm for ECM and 1mm for hair follicle composition. A 3d scatter plot shows the association of these two features with aging among all samples (**e**). Analysis of the occurrence of predictive paired features associated with cores of the aging process. In the top 100 most predictive bivariate pairs, the most frequently occurred pairs are between core 3 (hair and sebaceous gland shrinking) and core 10 (ECM alignment and Interstitial space widening) (**f**). **g.** The heatmap shows 99 tissue specimens (ranked in order by age) and the corresponding 108 age-associated features. Principal component analysis shows the first 29 principal components contain 95% of the total variance. A scatter plot shows the distribution of the sample along the first two principal components (PCs) and readily shows the chronological age of samples is associated with PC1 and PC2. **h,** Implementation of multivariate models with first 29 PCs for aging prediction. A violin plot shows the MAE of each sample from leave-one-out validation using GLM, Lasso, and SVM. A scatter plot shows predicted age (leave-one-out validation) vs chronological age using SVM in each sample. SVM achieved the lowest MAE of 7.7 years.

To explore the connection between aging-associated features, we primarily evaluated the prediction power of bivariate pairs among them, using the bivariate generalized linear regression (GLM). Our results showed that the MAE derived from a bivariate model was generally lower than the univariate MAE (**Figure 3d**). The best predictive bivariate pair was ECM alignment and hair follicle composition with an MAE of 11.7 years (**Figure 3e**). Among the top 100 most predictive bivariate pairs, 41 were pairs between core 3 (Hair and sebaceous gland shrinking) and core 10 (ECM matrix alignment and Interstitial space widening) (**Figure 3f, Supplementary Table 6**).

Finally, we investigated the age predictive power of these features using a multivariate model. Principal component analysis (PCA) was used to reduce the 108 features into 29 principal components (PCs), capturing 95% variance within the data set. The reduced PCs space readily exhibited a strong association with age (**Figure 3g**). We evaluated the predictive power of the reduced feature space with various machine learning (ML) algorithms including the generalized regression model (GLM), support vector machines (SVMs), Random Forest Regression (RF), and Lasso (least absolute shrinkage and selection operator), which drops some features by penalizing coefficients and driving them to zero (L2 penalty). The results showed that the SVM displayed the most accurate prediction with an MAE value of 7.7 years compared to 9.3 years from GLM and 8.9 years from Lasso. (**Figure 3h**, **Supplementary Table 7**).

Overall, our analysis indicates that morphometric features of the human skin are quantitatively linked to the aging process and could be further developed as biomarkers for aging.

### No significant effect of biological sex on aging in skin tissue

Next, we investigated whether the predictive power of aging-associated morphometric features of the skin was associated with sex. We found that the age predictive accuracy from the multivariate model was comparable between males and females, with an average MAE of 8.45 years for males and 7.76 years for females, respectively (**Figure 4a**). Analysis of sex-discrimination capability among 108 aging features, revealed seven features which had absolute effect size difference larger than 10% and p-value lower than 0.1 from two-tailed t-test (**Figure 4b** and **Supplementary Table 3**). Among the seven sex discriminant features, three were tissue/cell spatial distribution feature, three were dermal ECM fine structure features, and one was a tissue shape feature. We ranked the aging biomarkers based on the GLM accuracy in classifying the sex (**Figure 4c**). Stromal cell density was the top feature that differentiated biological sex with a classification accuracy of 66%, where samples from females had higher stromal cell density (169 N/mm^2^) than observed in samples from males (132 N/mm^2^) with a p-value of 0.78×10^−5^ (**Figure 4d**). Sex classification accuracy narrowly increased to 67% when using the multivariate GLM using the principal components reduced from the seven predictive features (**Figure 4e**). Overall, these results indicate that sex is not strongly associated with skin aging features of the back, as the inclusion of these features in the aging prediction does not significantly affect the prediction results (**Figure 4f**).

**Fig. 4:**
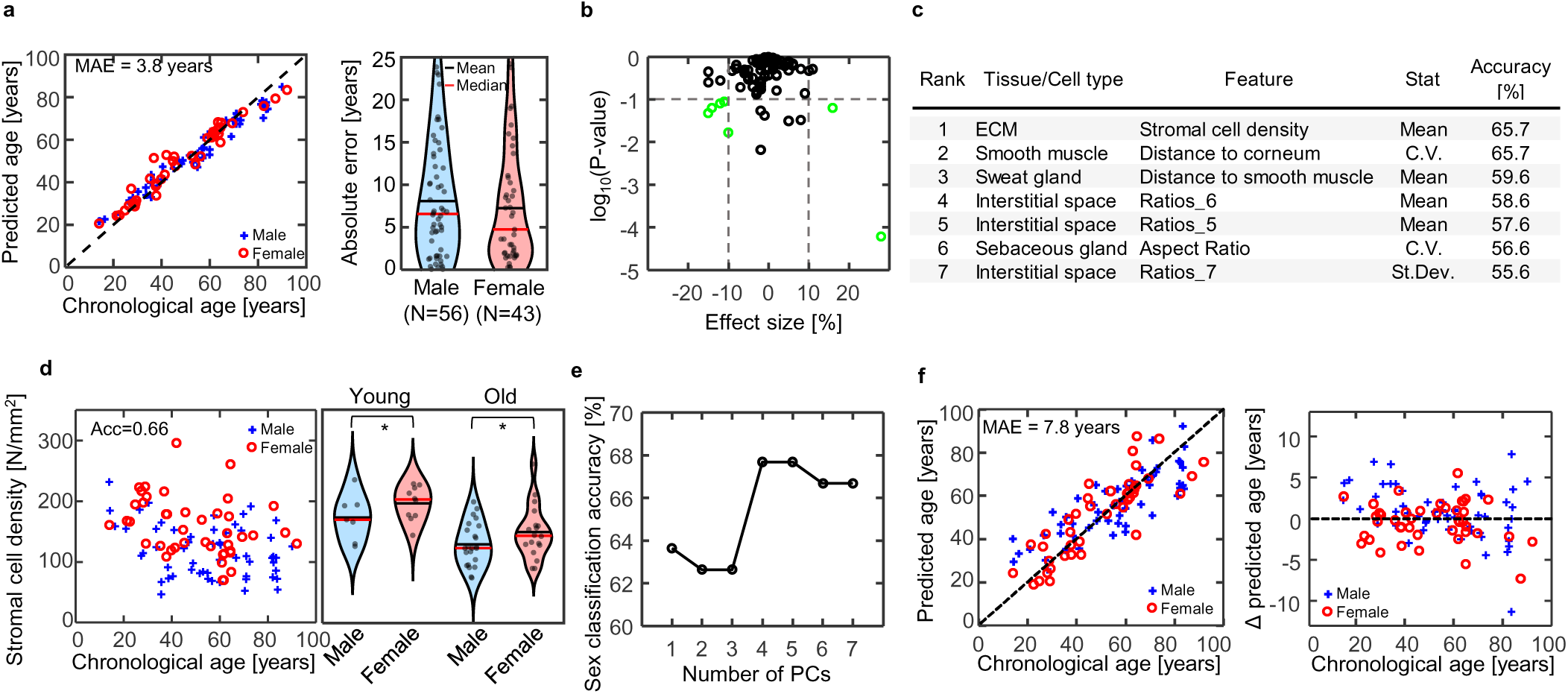
Skin aging features are not strongly associated with biological sex. **a.** A scatter plot of predicted ages and chronological ages in males and females. Predicted age is derived from leave-one-out using the multivariate SVM. MAE is 7.3 years among female samples compared to 8.1 years among male samples. **b.** A scatter plot illustrating the association between gender and 108 aging features with relative differences in mean and *p*-value (two-way t-test) between genders. Seven features have a relative difference of 10%+ with a p-value less than 0.1. **c.** Summary of the gender-associated features and their corresponding accuracy in classifying gender. **d.** A scatter plot shows the relation between stromal cell density and the age of donors in both genders. Violin plots comparing stromal cell density between genders in both young (30 years old or younger) and old (60 years old or older) show that stromal cell density is significantly higher in females than in males in both age ranges. **e.** The accuracy of gender classification is based on an increasing number of principal components derived from gender-associated features. The highest accuracy, reaching 68%, is attained using the first 4 principal components (PCs). **f.** Incorporating gender information into the aging prediction using the multivariate SVM model as depicted in Fig. 3h, a scatter plot illustrates the predicted age versus chronological age for donors of both genders. The Mean Absolute Error (MAE) is 7.8 years when gender information is considered in the prediction model. The scatter plot further depicts the difference in prediction errors for each sample, comparing models with and without considering gender. On average, there is a +0.8 years change in males and a −0.5 years change in females. The overall MAE increases by 0.1 year with the inclusion of gender information when compared to the MAEs obtained without considering sample gender information.

## Discussion

Using a deep learning-based framework, we present a rigorously curated and segmented atlas of normal skin microarchitecture at both tissue and cell levels from FFPE skin tissue sections. With the fully segmented skin tissue atlas, we captured the comprehensive cell and tissue morphometrics and associated spatial distributions. We identified ten core processes that occur in the aging skin. Among identified age-associated architecture changes, several features were consistent with previous studies, such as epidermis thinning^45–47^, hair loss^42,48^, decreased collagen content^49^, and collagen alignment and bundling^50^. Our study highlights five under-reported features of skin aging: 1) fewer and smaller pilosebaceous units; 2) narrower blood vessels; 3) increased distance between the blood vessel and epidermis; 4) Aligned ECM with epidermis; and 5) expanded interstitial space.

Our findings suggest that skin aging on the back is primarily characterized by progressive fibrosis and the loss of supporting structures in the dermis. This fibrosis is promoted by more rigid and aligned dermal extracellular matrix (ECM) where stromal cells are adapted into, likely resulting in the flattening and thinning of the epidermis and compressing interstitial structures such as hair follicles, sebaceous glands, blood vessels, and dermal adipose tissue (**Figure 5**). In addition, our results show that the decreased radius of blood vessels (**Figure 3b**) was one of the top predictors of aging and indicates the substantial vasculature architecture transformation as skin ages, which can, in part, explain the reduced blood circulations identified previously ^40,44,51^. Blood vessels play crucial roles in nutrient and oxygen delivery, waste removal, carbon dioxide exchange, and facilitating immune cell access ^52^. Furthermore, we observed that blood vessels tended to be located farther from the epidermis with increasing age (**Supplementary Figure 6**). These findings imply reduced nutrient delivery, contributing alongside dermal fibrosis to epidermal thinning (**Figure 2d**) due to slower basal cell replication rates ^52^. This vascular change could potentially drive the decrease in stromal cell density (**Figure 4d**) and ECM content, ultimately contributing to decreased dermal thickness and increased interstitial spaces as the dermis loses its structural integrity with age.

**Fig. 5:**
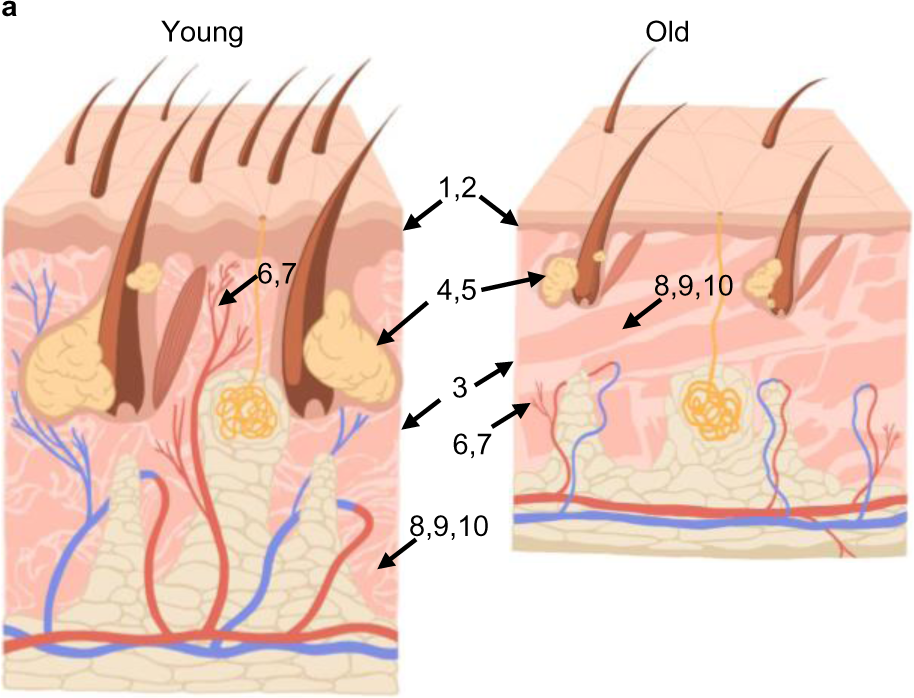
the effect of aging on the microanatomy of human skin. a. illustration of distinct macroscopic differences between young (age 10∼30) and old (age 60∼90) human skin, labeled with the corresponding number in the following list: 1 and 2. flatter and thinner epidermis, 3. thinner dermis, 4. fewer pilosebaceous units (a unit includes hair shaft, hair follicle, sebaceous gland, and arrector pili muscle), 5. smaller pilosebaceous units, 6. narrower blood vessels, 7. blood vessel distant from epidermis, 8. ECM more aligned with epidermis, 9. thicker ECM bundles, 10. wider interstitial bundles.

One primary mission in aging research is to identify intervention approaches that promote healthy aging and longevity by slowing down the aging process. Molecular analyses at the epigenetic, transcriptomic, and proteomic levels have provided potential molecular therapeutic targets for healthy aging^53^. These molecular analyses mainly reflect transformations at the cellular level^54,55^. Our tissue-level assessments further highlight that there is a large-scale coordinated transformation in tissue architectures with aging. The transformed tissue microanatomy could be closely associated with deteriorating tissue functions due to aging. Effective anti-aging therapy should reverse the course of tissue microanatomical transformation by editing the cells^56–58^. Therefore, evaluating the performance of aging drugs at the tissue microanatomy level can provide valuable insights and serve as an early surrogate marker for aging biomarkers^59^.

While all skin follows the same general architecture, significant variations occur throughout the body^60–63^. This study focused on evaluating samples from the human back, a large area of the body that suffers relatively less from body location variance^63^. However, other areas of the body show significantly different histologic features; scalp, for example, has an abundance of hair follicles and other dermal appendages^64,65^, whereas volar skin of the palms and soles of the feet has no hair, but presents a thickened epidermis and stratum corneum to withstand pressure^29^. Future directions include comparing the unique morphometric features of aging back skin with skin from other locations throughout the body. A combination of morphometric feature analysis with spatially resolved proteomics or genomic approaches may also provide significant insights on molecular drivers of age-related biophysical changes and changes within biological niches within the skin microenvironment. It is well established that image-based features derived from tissue images are often influenced by sample processing procedures, imaging parameters, and image quality ^66,67^. Therefore, developing an effective algorithm to normalize batch effects in cohorts with multiple batches is essential for further development and validation of generalized aging biomarkers in tissue sections.

## Methods

### Sample acquisition

Excisional skin specimens with substantial margin of normal tissue around the lesions were identified and collected from the archives at the Johns Hopkins Hospital. We compiled a cohort of patients whose skin samples were acquired as part of clinical routine from year 2015 to 2019 with underlying diseases, such as melanoma, keratosis, epidermal cyst, fibrosis, basal cell carcinoma, and squamous cell carcinoma. This cohort was compiled based on the volume of sample, evaluation on clinical notes whether the margin is free of disease, and their age distributions. The H&E stained tissue section of archive skin tissue samples from the cohorts were than obtained from the archive and imaged. The images of H&E stained sections were then evaluated by a board-certified dermatopathologist to identify the sections with grossly normal skin margins free of solar elastosis, inflammation, cicatrix, and the primary underlying disease.

Each slide contains between 1 and 6 tissue sections, depending on the size of the biopsy. Larger biopsies are split and placed into the same tissue cassette to form a formalin-fixed paraffin embedded (FFPE) block. All tissue sections on a slide are evaluated, and only the sections free of diseases are used for further analysis. Out of 295 patient samples, 165 samples were selected. Among these samples, the majority (N=130) of tissue blocks are from the skin from the back, and the rest are from the abdomen, chest, head, neck, arms, and legs. Out of 130 back tissues, 101 are from Caucasian patients.

### Image acquisition

Each selected H&E tissue slide is scanned at 20x magnification using a Hamamatsu Nanozoomer S210 scanner with a resolution of 0.454 mm per pixel. Nine focus points are used per tissue section.

### Tissue semantic segmentation

To segment skin tissue objects (e.g., glands, smooth muscles, etc.), we used a DeepLabV3+ CNN architecture^68^ pretrained on ImageNet^69^. To train the CNN for tissue labeling, we first establish the training data set. We randomly selected 20 out of the above 165 tissue images and manually annotated them using Aperio ImageScope. We labeled twelve tissue components in the skin including: 1. stratum corneum, 2. stratum spinosum, 3. hair shaft, 4. hair follicle, 5. Smooth muscle, 6. sebaceous gland, 7. sweat gland, 8. Nerve, 9. blood vessel, 10. ECM, 11. fat, and 12. interstitial space. The annotated tissue components are validated by a board-certified dermal pathologist. The DeepLabV3+ model was trained, cross-validated, and applied to all 165 whole slide images for pixel-wise classification of these 12 tissue components. Our skin tissue segmentation is validated using 5 independent, randomly selected samples. The model’s precision is 96.61%, and its recall is 93.94% (**Supplementary Figure 3**).

### Rotational alignment of skin tissue

The epidermis was labeled by a deep learning model. The labeled epidermal is converted to 2D point clouds^70,71^ (**Supplementary Figure 6**). The Y coordinate of the point clouds is subtracted by their means in Y-axis direction to be flattened along the horizontal X-axis. These flattened point clouds are the target for the alignment. Next, we compute the rotational angle needed to minimize the error between the original point clouds and the target using singular value decomposition. This type of problem is called an orthogonal Procrustes problem^72^. Once the rotational angle is found, we can use it to rotate each skin tissue section so that the horizontal axis of the resulting image is aligned with the skin surface. Detailed mathematical method is the following. Given that original point cloud *Q* = {*q*_1_, …, *q*_*N*_} and target cloud *P* = {*p*_1_, …, *p*_*N*_} has correspondences *C* = {(*i*, *j*)}, the error between the two are defined as *E*(*R*, *t*) = ∑_(*i*,*j*)∈*C*_ ‖*q*_*i*_ − *Rp*_*j*_ − *t*‖^2^. Singular value decomposition (SVD) is used to decompose the cross-covariance matrix *W* = ∑_(*i*,*j*)∈*C*_ *q*_*i*_′ *p*_*j*_′^*T*^ to *W* = UDV^*T*^, providing the values of U, D, V^T^. Then, the rotational matrix can be found as *R* = UV^*T*^. This matrix is 2×2 dimensions with the following elements, 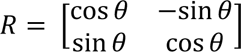. Finally, we take arctan of 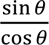 from the rotational matrix to compute the rotational angle. When the transpose of the rotational matrix has negative determinant, the matrix is in a reflection position, resulting in a negative value of rotational angle. To compute the correct angle, we multiply −1 to the rotational angle to flip its sign.

### Tissue composition and architecture quantification

The fully labeled skin tissue images allowed us to assess the composition of tissue subtypes and their respective organization. The composition is calculated by counting the number of pixels occupied by each tissue component and dividing it by the total number of pixels in the whole tissue section. The tissue architectural features such as thickness, waviness, roughness, etc. A detailed list of tissue architecture features can be found in the **supplementary table 2**.

### Nuclei detection

A deep neural network model was applied to detect hematoxylin-stained nuclei. HoVerNets, a convolutional neural network for nuclear segmentation in H&E^73^, was trained using several images selected arbitrarily from each group of tissue component for a total of N samples. Detailed workflow for training the model is illustrated in **Supplementary Figure 3**. Our implementation of HoVerNet takes a different approach from the original workflow in training the model to associate the Human Skin H&E images with nuclear regions. After training for 100 epochs, the HoVerNet was applied to detect nuclei for all 165 whole slide images, producing output images of predicted nuclei regions along with the coordinates of nuclei contour. Each tissue section size from 15mm^2^ ∼ 35mm^2^ generated 5000∼12000 detected nuclei. From the entire Skin tissue dataset, we detected 7.8 million nuclei. To determine the cell type of the nuclei, we use the location of nuclei centroid with regards to the tissue component label to identify which tissue component is a nucleus reside in. For instance, a nucleus detected within sweat gland region is almost certainly from a sweat gland cell. The spatial correlation between neighboring nuclei is quantified using metrics such as distance between neighboring cells using K-nearest neighbor algorithm (KNN), alignment between adject cells, and local nuclei population density within a tissue component object.

### Nuclear morphology

The nuclei contours are used to quantify morphological features of the nuclei, including, but not limited to, area, circularity, aspect ratio, orientation, and extent. A detailed list of nuclear morphology features can be found in the **Supplementary Table 2**. The area A is the total number of pixels within the contour and then converted to micrometer squared given that width and height of pixel are both 0.454µm. Perimeter P is defined by the length of contour line. The circularity is computed using the equation: 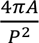. The aspect ratio is defined as the major axis length divided by minor axis length. The orientation is quantified by a conic fitting algorithm described by Fitzgibbon et al^74^. The extent is defined as area of contour divided by area of bounding box.

### Software

The following software tools were used for the predictor and related analysis: python 3.8.10, tensorflow-gpu 2.4.1, tensorflow-probability 0.11.1, scipy 1.6.2, statsodels 0.12.0, scikit-learn 0.23.2, pandas 1.1.1, seaborn 0.11.1 and numpy 1.21.6, R 4.0.2, MATLAB R2024a.

### Statistics and reproducibility

A correlation between a sample’s age and a morphometric feature was made using spearman rank coefficient. The linear regression p value for each morphometric feature tests the null hypothesis that the feature has no correlation with a sample’s age.

## Acknowledgements

This work was supported through grants from the National Cancer Institute (U54CA143868 and U54CA268083 to DW, PW), National Cancer Institute (UG3CA275681 to PW, DW) the National Institute of Arthritis and Musculoskeletal and Skin Diseases (U54AR081774 to DW) and the National Institute on Aging (U01AG060903 to DW, JMP), This study was supported by the Johns Hopkins Older Americans Independence Center National Institute on Aging (P30 AG021334 to JW, JMP)

## Contributions

KSH, IS, CJ, PW, DW, JMP conceived, developed, and designed the study. KSH, ER, CB performed sample collection. KSH, IS, JS developed skin tissue model. KSH, BS, AK, PW, DW wrote and edited the manuscript with input from all co-authors. PW, DW, JMP supervised study and secured funding.

---

**Table 1:** Top 10 univariate features.

| Rank | Feature ID | Tissue component | Feature name | Feature type | Core process | MAE [years] |
| --- | --- | --- | --- | --- | --- | --- |
| 1 | 218 | Blood vessel | Mean bounding box height | 4 | 6 | 13.1 |
| 2 | 116 | Hair follicle | Content per length scale | 4 | 5 | 13.5 |
| 3 | 4 | Hair follicle | Composition | 1 | 5 | 13.5 |
| 4 | 76 | S. Spinosum | Content per length scale | 4 | 6 | 14.0 |
| 5 | 121 | Hair follicle | Occurrence per length scale | 4 | 5 | 14.0 |
| 6 | 27 | ECM | Anisotropy_B | 3 | 1 | 14.0 |
| 7 | 29 | ECM | Anisotropy_B_masked | 3 | 1 | 14.0 |
| 8 | 160 | Sebaceous gland | Bounding box height st.dev. | 4 | 5 | 14.1 |
| 9 | 153 | Sebaceous gland | Size st.dev. | 4 | 5 | 14.1 |
| 10 | 163 | Sebaceous gland | Major axis length st.dev. | 4 | 5 | 14.2 |

**Table 2:** Top 10 bivariate features.

| Rank | Feature ID | Tissue component | Feature name | Feature type | Core process | MAE [years] |
| --- | --- | --- | --- | --- | --- | --- |
| 1 | 4 | Hair follicle | Composition | 1 | 5 | 11.7 |
|  | 29 | ECM | Anisotropy_B_masked | 3 | 1 |  |
| 2 | 4 | Hair follicle | Content per length scale | 1 | 5 | 11.7 |
|  | 27 | ECM | Anisotropy_B | 3 | 1 |  |
| 3 | 160 | Sebaceous gland | std bounding box height | 4 | 5 | 11.8 |
|  | 218 | Blood vessel | mean bounding box height | 4 | 6 |  |
| 4 | 29 | ECM | anisotropy_B_masked | 3 | 1 | 11.8 |
|  | 116 | Hair follicle | content per length scale | 4 | 5 |  |
| 5 | 29 | ECM | anisotropy_B_masked | 3 | 1 | 11.8 |
|  | 218 | Blood vessel | mean bounding box height | 4 | 6 |  |
| 6 | 171 | Sebaceous gland | std perimeter | 4 | 5 | 11.8 |
|  | 218 | Blood vessel | mean bounding box height | 4 | 6 |  |
| 7 | 27 | ECM | anisotropy_B | 3 | 1 | 11.8 |
|  | 116 | Hair follicle | content per length scale | 4 | 5 |  |
| 8 | 153 | Sebaceous gland | std size | 4 | 5 | 11.9 |
|  | 218 | Blood vessel | mean bounding box height | 4 | 6 |  |
| 9 | 27 | ECM | anisotropy_B | 3 | 1 | 11.9 |
|  | 218 | Blood vessel | mean bounding box height | 4 | 6 |  |
| 10 | 153 | ECM | anisotropy_A | 3 | 1 | 11.9 |
|  | 218 | Blood vessel | mean bounding box height | 4 | 6 |  |

