## Supplemental Figure for "qMAP enabled microanatomical mapping of human skin aging"

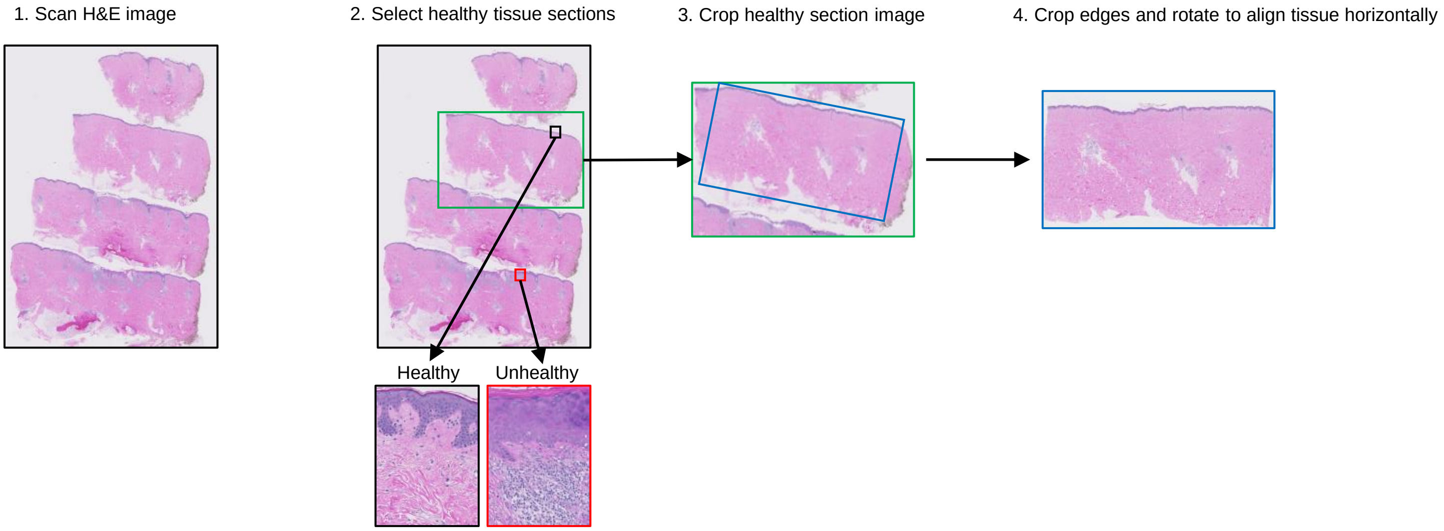


**Fig. S1. Detailed procedures of figure 1a "H&E imaging & healthy tissue screening"**

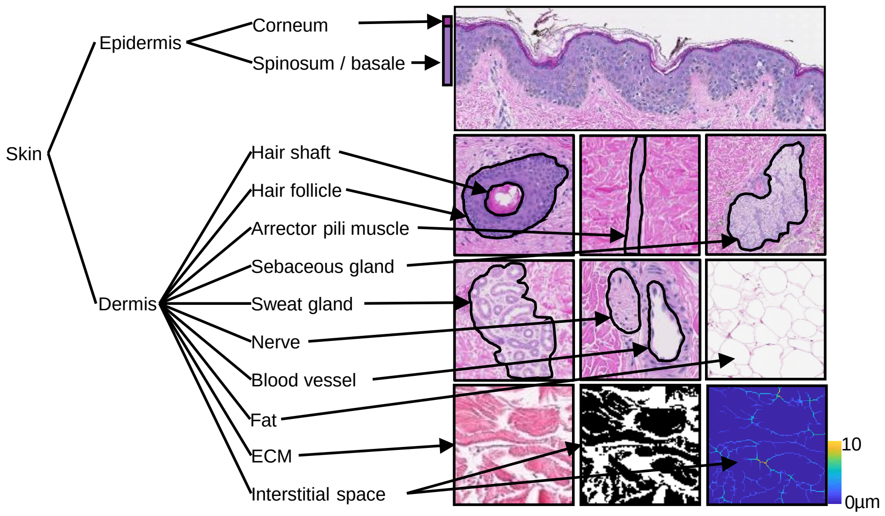


**Fig. S2. Tissue component examples**

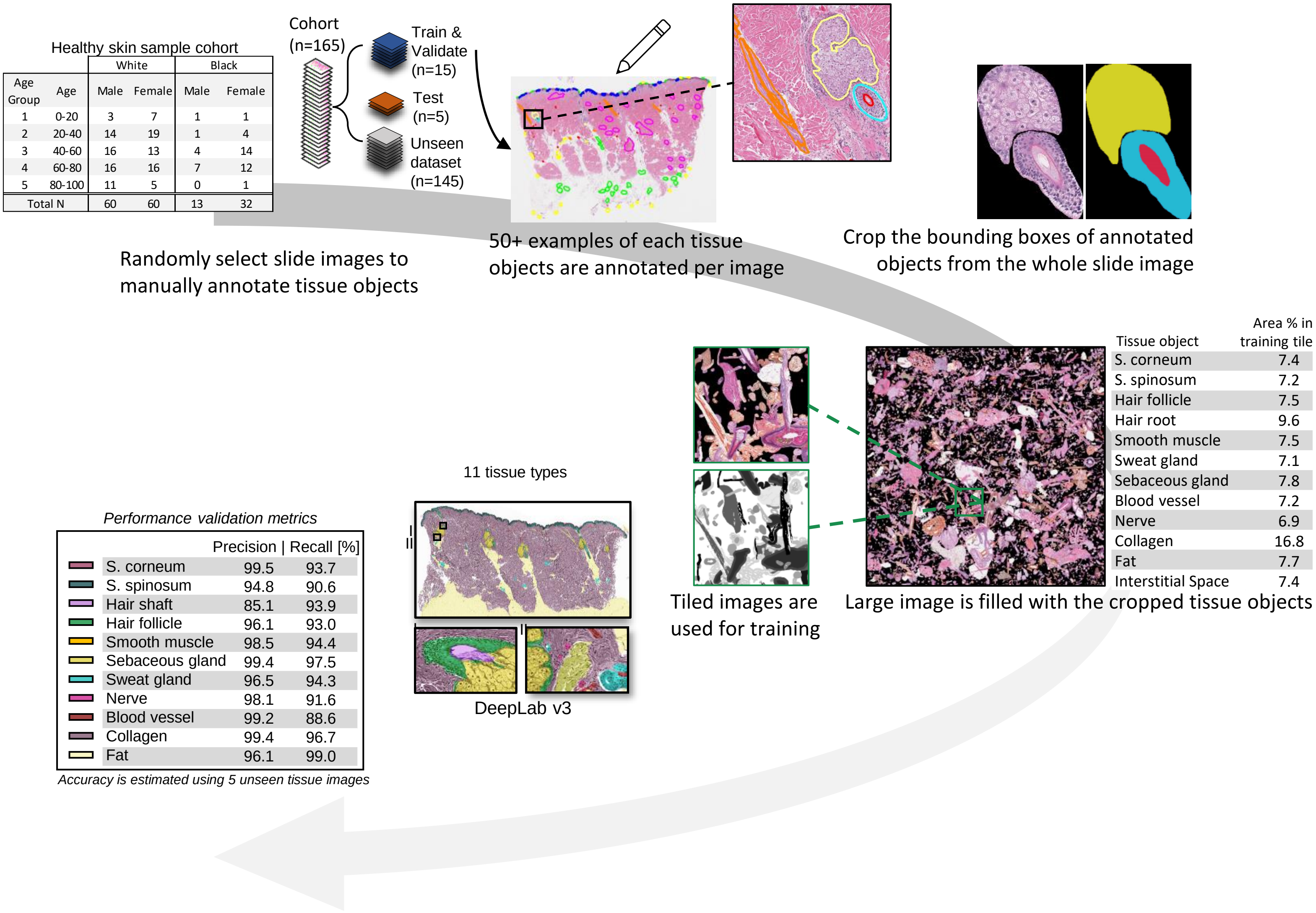


**Fig. S3. Overview of semantic segmentation model training.** (A) 20 out of 109 samples were randomly selected for manual annotation. Minimum 50 examples of each tissue object were annotated. The annotated objects are cropped from the whole slide image. (B) To construct training and validation sets, cropped annotations were overlayed on a 9000x9000 large image until the image was >65% full and such that the number of annotations of each type was roughly equal. While overlaying, the cropped annotations are augmented individually via random rotation, scaling by a random factor between 0.8 and 1.2, and hue augmentation by a factor of 0.8-1.2 in each RGB color channel. (C) The constructed large images of overlaid annotations were tiled into smaller size of 500x500 for training and validation. Additional large images are prepared the same way from 5 independent, randomly selected samples for testing.

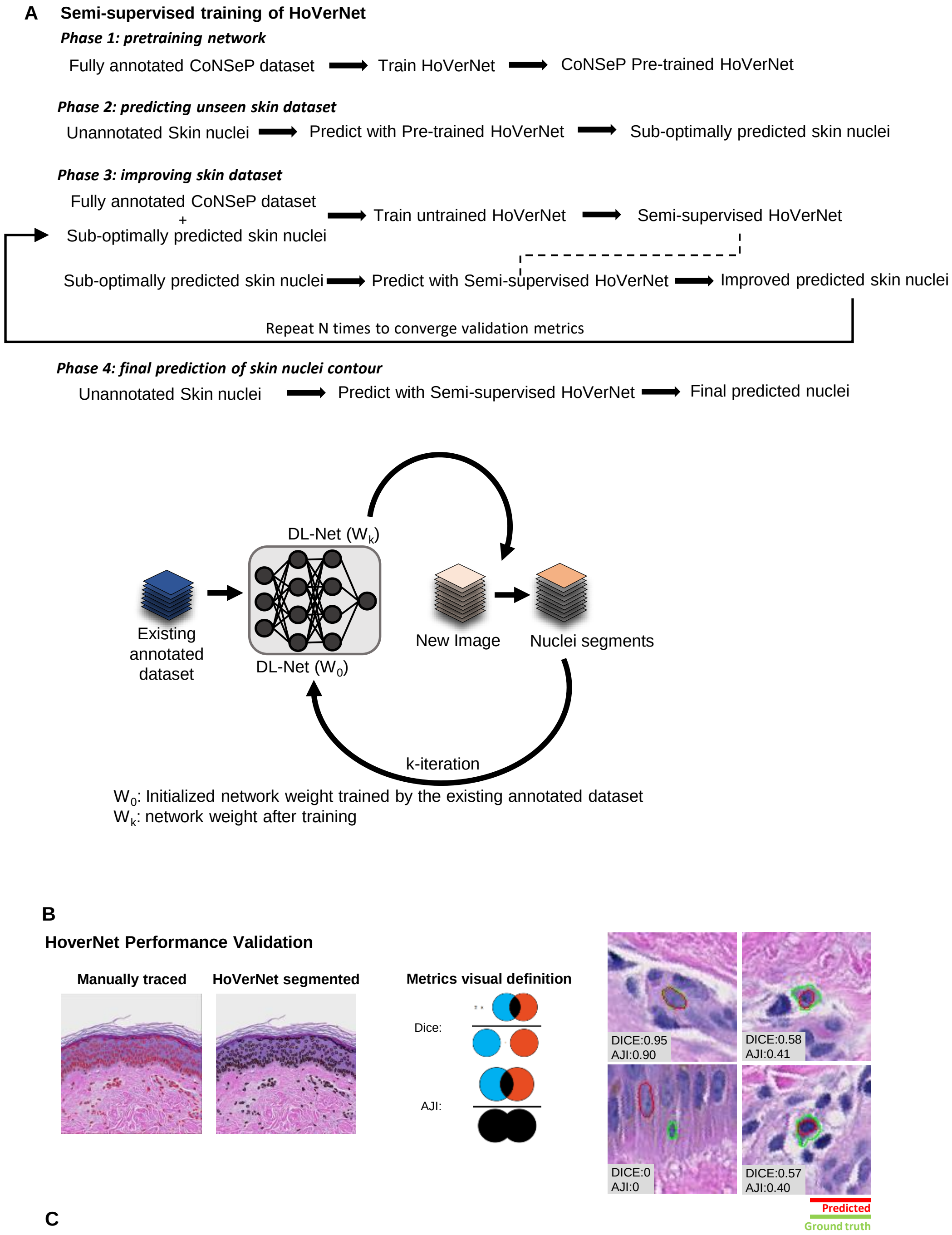


**C**

| Validation Methods | DICE | AJI | DQ | SQ | PQ |
|---|---|---|---|---|---|
| ImageNet | | | | | |
| CoNSeP | 0.66 | 0.40 | 0.53 | 0.76 | 0.41 |
| CoNSeP + Skin Iteration 1 | 0.70 | 0.79 | 0.64 | 0.77 | 0.80 |
| CoNSeP + Skin Iteration 2 | 0.71 | 0.85 | 0.64 | 0.76 | 0.87 |
| CoNSeP + Skin Iteration 3 | 0.71 | 0.88 | 0.65 | 0.78 | 0.87 |
| CoNSeP + Skin Iteration 4 | 0.73 | 0.91 | 0.70 | 0.81 | 0.88 |
| CoNSeP + Skin Iteration 5 | 0.73 | 0.90 | 0.72 | 0.80 | 0.90 |

CoNSeP dataset contains 26 and 14 train and test set with 1000x1000 pixel images
Skin dataset contains 787 and 140 train and test set with 1000x1000 pixel images

**Fig. S4. Evaluation of instance segmentation models with CoNSeP dataset and our custom dataset.**

**A** ***Tissue composition (12 features)***

| Age | Young 0-30 | Middle-aged 30-60 | Old 60-100 | Average 0-100 |
|---|---|---|---|---|
| Nerve | 0.4 | 0.3 | 0.3 | 0.3 |
| Corneum | 0.6 | 0.7 | 0.8 | 0.7 |
| Hair shaft | 0.1 | 0.1 | 0.1 | 0.1 |
| Sebaceous gland | 1.0 | 0.7 | 0.3 | 0.6 |
| Hair follicle | 0.4 | 0.2 | 0.1 | 0.2 |
| Spinosum | 2.2 | 1.9 | 1.8 | 1.9 |
| Blood vessel | 0.9 | 1.0 | 0.9 | 0.9 |
| ECM | 81.0 | 78.1 | 79.7 | 79.2 |
| Sweat gland | 0.6 | 0.7 | 0.5 | 0.6 |
| Arrector pili muscle | 0.6 | 0.6 | 0.5 | 0.6 |
| Fat | 3.8 | 4.6 | 4.9 | 4.6 |
| Interstitial Space | 7.8 | 11.1 | 9.9 | 10.0 |

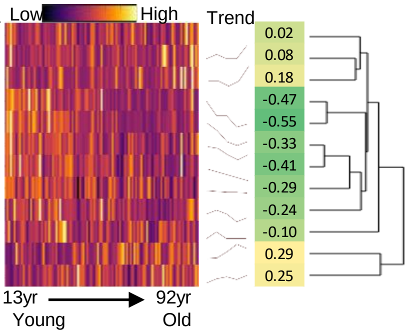


### Fig. S5. Change in skin tissue composition
